# A mitochondrion-like ribosome in an insect endosymbiont

**DOI:** 10.64898/2026.10.18.759926

**Authors:** Israel S. Fernández, Jessica M. Warren, Justus Nwachukwu, Dewight Williams, Amaia González-Magaña, Dalton Leprich, Noa Spencer, John P. McCutcheon

## Abstract

Understanding how the mitochondrion evolved from a bacterial endosymbiont to an organelle is obscured by time and a lack of comparative examples. It is clear that mitochondrial ribosomes are derived from bacterial ribosomes, but it is unclear whether the baroque and idiosyncratic structural changes seen in mitochondrial ribosomes are specifically driven by mitochondrial evolution or simply reflect the syndrome of being a long-term endosymbiont. Here we show that three characteristic features of mitochondrial ribosomes— loss of ribosomal proteins of bacterial origin, acquisition of eukaryotic host proteins, and alterations to specific structural regions—are shared by *Hodgkinia*, an ancient bacterial endosymbiont of cicadas. These results reveal how endosymbiont ribosomes evolve in response to the loss of numerous ribosomal proteins, and imply that many of the striking changes observed in mitochondrial ribosomes result from their ancestry as bacterial endosymbionts rather than their uniqueness as mitochondria.

## INTRODUCTION

The bacterial ribosome is an extraordinarily conserved machine. The presence of 3 RNAs and 54 proteins in most extant bacterial ribosomes imply that this modern structure was found in the first bacterial cell more than 3 billion years ago ^1,2^. This strong conservation across bacteria also suggests that the ribosome in the alphaproteobacterial ancestor of the mitochondrion had a canonical bacterial structure ^1,3,4^, and that the substantial structural changes observed in mitochondrial ribosomes (mitoribosomes) ^5–8^ have occurred during or after the establishment of the organelle. Mitoribosomes have lost many of their ancestral bacterial ribosomal proteins (bRPs), contracted or expanded their ribosomal RNA (rRNA), extended bRP lengths compared to their homologs in bacteria, and acquired numerous new proteins of eukaryotic ancestry that are imported from the host (mRPs) ^9^. Two regions of the mitoribosomal large subunit (LSU) are repeatedly found to be altered compared to bacterial ribosomes: the central protuberance (CP) and the nascent polypeptide exit tunnel (ET). The 5S rRNA comprises the core of the CP, and in mitoribosomes the 5S rRNA has been reduced and modified in green algae ^7,10^, lost and replaced by a tRNA in animals ^11^, and lost and replaced by numerous new eukaryotic proteins in trypanosomes ^12^. Similarly, in yeast mitoribosomes, new eukaryotic proteins have been recruited to the ET which, in combination with expansions in RPs and rRNA, have moved the ET about 35 Å away from its conserved location in bacteria ^13^. The ET in trypanosome mitoribosomes has also undergone significant structural change, with an alternative ET path formed by protein extensions of bRPs and the recruitment of new mRPs ^12^. It is unclear whether these dramatic changes in mitoribosomes reflect specific requirements of mitochondrial biology or are more general outcomes of a long endosymbiotic relationship with eukaryotic cells.

Many cellular and genetic features once thought to be unique to mitochondria and plastids have now been identified in bacterial endosymbionts of insects and protists ^14,15^. Extreme genome reduction ^16^, biochemical and genomic complementarity ^17,18^, and import of host proteins into endosymbiont cells ^17,19,20^ have been described for several systems containing stably associated endosymbiotic bacteria. Endosymbionts tend to lose more genes the longer they spend in a host-restricted state ^21^. Mirroring the deep evolutionary conservation of the bacterial ribosome, genes involved in translation are among the last to be lost as genomes shrink ^15,22–24^. Some of the most reduced endosymbiotic genomes are predicted to have lost multiple tRNAs and aminoacyl-tRNA synthetase genes, along with up to 20 of the 54 bRPs ^15,21,23–25^. The alphaproteobacterial cicada endosymbiont *Candidatus* Hodgkinia cicadicola (hereafter *Hodgkinia*) is one such example. Its tiny 144 kb genome is predicted to have lost genes for several tRNAs and aminoacyl-tRNA synthetases ^26^, along with 13 bRPs ^25^. It is unclear how endosymbionts such as *Hodgkinia* evolves in response to such extensive gene loss in these almost universally conserved translation-related processes. Here, using a combination of genomics, transcriptomics, and cryo-electron microscopy (cryo-EM), we show that the *Hodgkinia* ribosome has undergone changes that strongly parallel mitoribosome evolution despite having an independent evolutionary origin.

## RESULTS

### Preparation of ribosomes from cicada endosymbionts and structure determination

In the summer of 2025 we captured 2 male *Diceroprocta semicincta* cicadas in the Phoenix area and dissected out their bacteriomes, the specialized tissue containing *Hodgkinia* and its co-symbiont, *Candidatus* Karelsulcia muelleri (hereafter *Karelsulcia*) (**Figure 1A**) ^27^. Cicada bacteriome tissue contains a mix of ribosomes from four sources: the cicada cytosol, the cicada mitochondria, and ribosomes from both endosymbionts, *Hodgkinia* and *Karelsulcia*. We miniaturized the RAPPL protocol ^28^ to purify ribosomes from small amounts of bacteriome tissue. This approach produced enough material to plunge-freeze four cryo-EM grids from dissections of our two cicadas. One of these grids was of sufficient quality in terms of ice thickness and particle distribution for data collection (**Figure 1A**).

**Figure 1.**
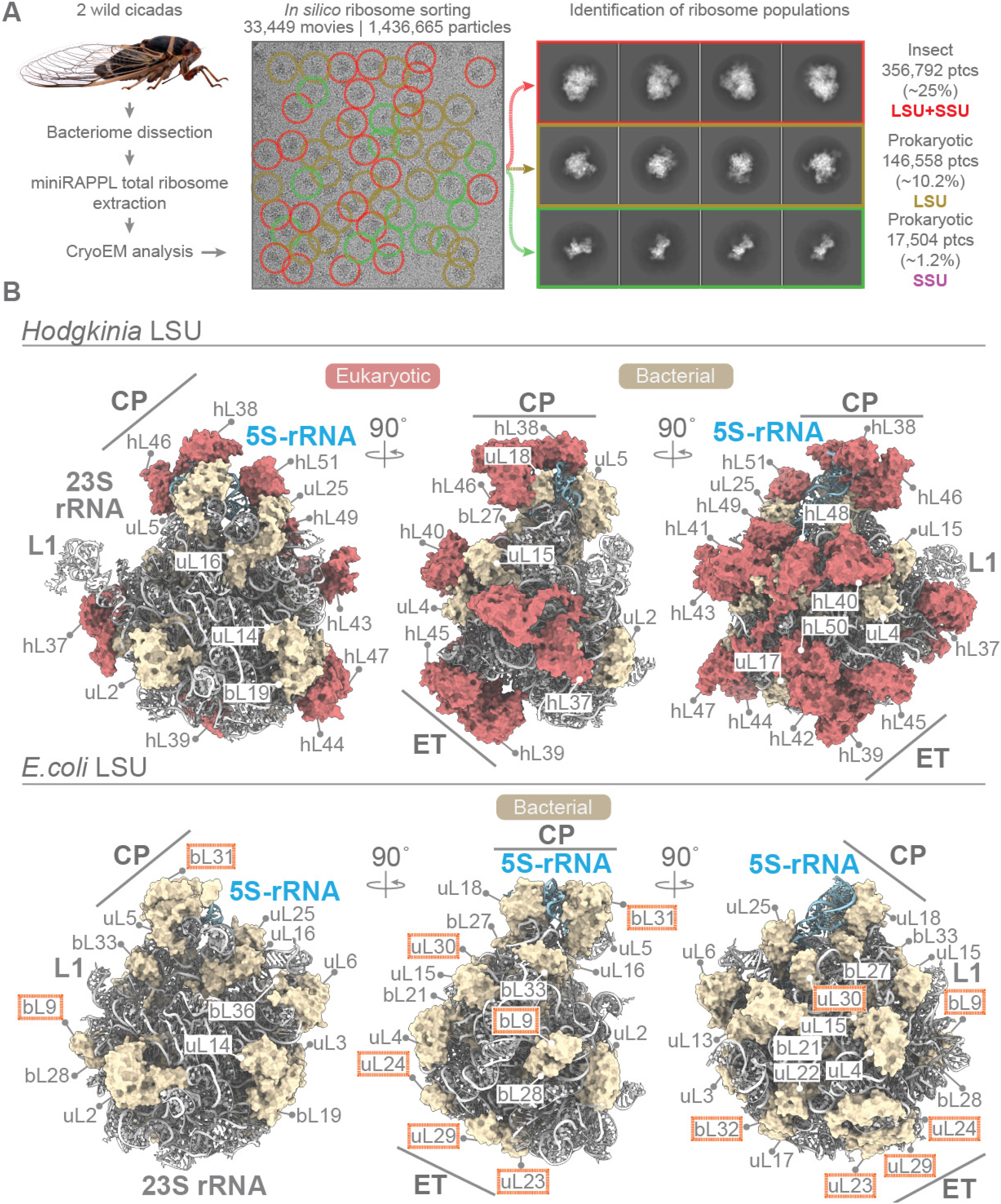
Overview of the experimental strategy and the *Hodgkinia* LSU structure compared to *E. coli*. (**A**) Schematics of sample preparation (left) and cryo-EM analysis showing a representative motion corrected image (center) with ribosomes labeled according to their origin, with eukaryotic ribosomes from the cicada host in red, bacterial large subunits in yellow and bacterial small subunits in green. The final reference-free 2D class averages including information from all particles on the dataset are shown on the right side of (A). (**B**) The top part of the figure shows orthogonal views of the *Hodgkinia* LSU with 23S rRNA depicted as grey cartoons, 5S rRNA as blue cartoons, ribosomal proteins of bacterial ancestry as yellow molecular surfaces, and new ribosomal proteins from the cicada genome in surface representation colored red. For comparison, the bottom part of the figure shows corresponding orthogonal views of the *E. coli* LSU (PDB ID 7k00) with the same visualization settings. *E. coli* ribosomal proteins framed in orange boxes are those missing from the *Hodgkinia* LSU and genome.

As expected, *in silico* ribosome sorting using empirical Bayesian methods ^29^ revealed a mixture of ribosome populations. While fully assembled host cytosolic complexes (80S ribosomes) dominated the dataset, minor populations of bacterial-like classes were visible in the 2D class-averages (**Fig. 1A**). These classes exhibited features in length, diameter and overall shape reminiscent of the bacterial LSU (50S in bacteria) and the bacterial small subunit (SSU, 30S in bacteria). Fully assembled bacterial-like ribosomes (70S) were not identified in the dataset.

The 2D classes corresponding to a bacterial-like LSU (~146,500 particles) produced a 3D map with nominal overall resolution of ~3 Å (**Figures 1B** and **S1, Table S1**). We also identified a bacterial-like SSU approximately one order of magnitude less abundant than bacterial-like LSU (~17,500 particles). The reduced number of particles combined with severe preferential orientation on the grid limited the attainable resolution in the 3D reconstruction of the SSU to ~12 Å (**Figure S2**). We did not pursue further analysis of this SSU given its low resolution.

### Predicted losses of bRPs and the identification of the LSU map

Because the cicada species in this study has two bacterial endosymbionts, *Hodgkinia* and *Karelsulcia*, along with its own mitochondria, the reconstructed bacterial LSU particle could have originated from either bacterium, the mitochondrion, or a mixture of the three. We rejected the mixture hypothesis given the quality of the map in terms of resolution and homogeneity (**Figure S1**), and we rejected the mitochondrial hypothesis due to the size of the particle. In order to identify whether this LSU reconstruction belonged to *Hodgkinia* or *Karelsulcia*, we followed a two-pronged approach rooted in the extreme conservation of the ribosomal peptidyl-transferase center (PTC) along with the predicted differences in the sets of ribosomal proteins in the *Hodgkinia* and *Karelsulcia* genomes ^25^. We first fitted an atomic model of the LSU rRNA from *E. coli* ^30^ into our map, which allowed an unambiguous and precise fitting of the *E. coli* PTC. Using the *E. coli* PTC as a structural anchor, we superposed the rest of the *E. coli* LSU onto our map. Most *E. co* ribosomal proteins fit the density of the map well, but several had no corresponding map density, even at low contour values, suggesting the loss of multiple bRPs from this LSU (**Figure S3B**).

Genome annotation for *Karelsulcia* predicts that only uL29 and uL30 are missing from its LSU compared to *E. coli*, while *Hodgkinia* is predicted to be missing 10 bRPs from the LSU (uL1, bL9, bL21, uL23, uL24, uL29, uL30, bL31, bL32, bL34) ^25^. Our preliminary structural analysis revealed weak or nonexistent density at 9 of the 10 bRPs predicted to be missing in *Hodgkinia* (**Figure S3B**). These structural patterns suggested that this LSU map was from *Hodgkinia*, not *Karelsulcia*, which we further verified using characteristic and discriminating amino acid residues in conserved bRPs. We observed robust map density corresponding to bL21 in *E. coli*, which was the sole conflict between our cryo-EM and genomic data. Re-analysis of the *Hodgkinia* genome revealed that the prediction of the bL21 gene as missing in the genome ^31^ was due to an annotation error. Overall, these results confirm that this map is from *Hodgkinia*, and that its LSU is missing 9 bRPs. Of note are the losses of uL23, uL24, and uL29, which are retained even across the diversity of mitoribosomes ^1^.

### The *Hodgkinia* LSU structure is a mosaic of bacterial and eukaryotic proteins

After confirming that our LSU map was from *Hodgkinia*, we built and refined the majority of the LSU 23S rRNA, the complete 5S rRNA, and 21 of the 24 *Hodgkinia* bRPs (the three stalk proteins uL10, uL11, and bL12 were not well ordered in our map (**Figure S4**), consistent with other LSU structures ^32^). After the removal of all map regions where we could find direct one-to-one mapping to homologous structures in *E. coli*, the difference map revealed substantial additional densities. These densities were located at the periphery of the LSU, with several clusters of density at the CP and ET (**Figure S3A**). In order to identify these unexpected regions of density, we used ModelAngelo ^33^ to generate structure-based profile hidden Markov models (HMMs) from our *Hodgkinia* LSU map and searched these HMMs against predicted proteomes from the *Hodgkinia* and *Karelsulcia* genomes ^27,31^ along with a set of host proteins predicted from a previous cicada transcriptome study ^26^ using the HMMER ^34^ function in ModelAngelo.

Outside of the expected *Hodgkinia* bRPs, we found no other high confidence HMM hits in predicted *Hodgkinia* or *Karelsulcia* proteomes. Remarkably, we found numerous high-confidence HMM hits to proteins predicted from our cicada transcriptome ^26^. However, because this transcriptome was produced using short reads and we had no reference cicada genome, the overall quality of the transcriptome was low and we were unable to disambiguate true splice variants from transcriptome misassembly. We generated a PacBio cicada genome (4.9 Gb; 1,396 contigs; N50 of 13.2 Mb; ~18X coverage) and a PacBio IsoSeq cicada transcriptome (22.8 M reads; avg. length 1,487 nts) to provide definitive references for our ModelAngelo structure-based search. These genomic and transcriptomic data confirmed our high-confidence HMM hits to predicted cicada proteins, and allowed us to unambiguously align these protein-producing transcripts onto the cicada nuclear genome (**Figure S5**). Using a combination of AlphaFold 3 ^35^ predictions and manual building and refinement, we placed 15 nuclear-encoded cicada proteins into the regions of unexplained density on the *Hodgkinia* LSU map with high confidence (**Figure 1B, Table S2**). Following the naming convention used for mitoribosomes, we have named these proteins hL37-hL51. The ‘h’ prefix reflects the presence of these proteins on the *Hodgkinia* ribosome, and the numbering starting at 37 reflects our assumption that the ribosome of *Hodgkinia*’s alphaproteobacterial ancestor had the complete set of bacterial LSU bRPs, the numbering of which ends at bL36 ^36^. We note that this set of 15 host proteins may be an undercount, as some regions of the *Hodgkinia* LSU map show additional density at low contour levels (**Figure S4**). We were unable to identify additional proteins in these regions due to their flexible and/or heterogeneous nature.

### The 15 host proteins are of various fold types and predicted functions

Unlike mRPs of eukaryotic ancestry that have accumulated on mitoribosomes, which tend to be from related families of alpha-helical rich proteins such as pentatricopeptide repeat proteins ^9,10^, the cicada proteins we find in the *Hodgkinia* LSU structure (hRPs) are a diverse mixture of fold types and have a variety of predicted functions seemingly unrelated to ribosome biology (**Table S2**). All but one cicada hRP have predicted N-terminal signal peptides (**Table S2**), suggesting that these proteins may be trafficked through the cicada secretory pathway as part of their targeting to *Hodgkinia* cells.

Two of these proteins, hL44 and hL47, are alternatively spliced versions of the same gene that physically interact with each other on the *Hodgkinia* LSU (**Figures 2** and **S6**). The genomic locus encoding these two splice variants is complex, with seven different splice variants all sharing the same first exon but different second exons, which apparently produce six different but related beta-1,3-glucan binding proteins (**Figure S6**). The introns of this locus are large, with the second exons of hL47 and hL44 located 154 kb and 158 kb downstream from their shared exon 1, respectively. These intron lengths are on the same order as the largest intron reported in *Drosophila* (268 kb), and far exceed the *Drosophila* average length 73 bp ^37^. hL47 interacts with bacterial ribosomal protein uL3, while hL44 interacts with both uL3 and bL19 (**Figure 2A**). A C-terminal loop of hL47 (encoded by its unique exon 2’) extends into a pocket of hL44 (encoded primarily by the shared exon 1) and makes extensive hydrophobic and complementary charge interactions with its neighboring protein isoform (**Figure 2B**).

**Figure 2.**
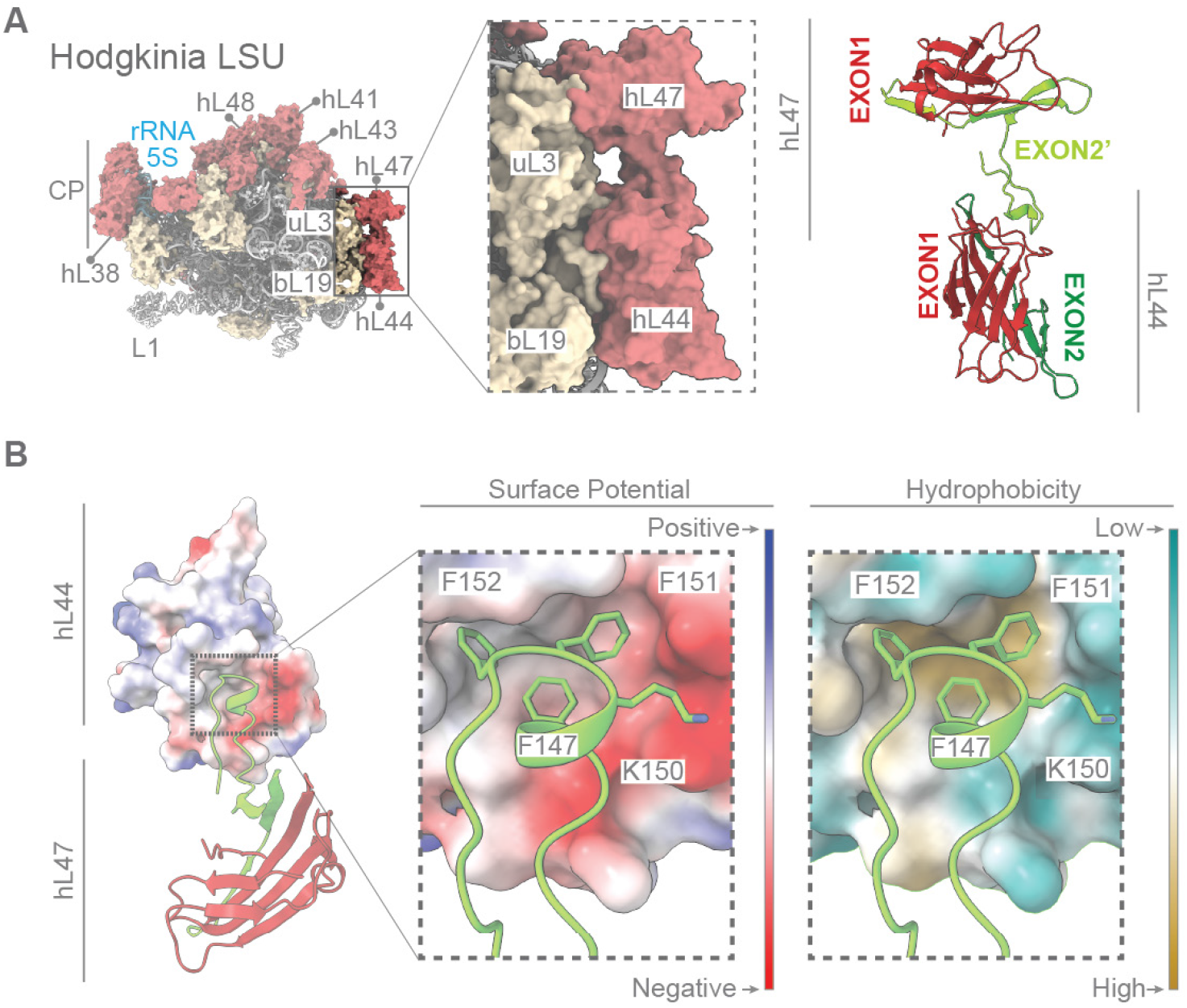
Interactions between hL44 and hL47, two cicada proteins resulting from alternative splicing of a single cicada gene. **(A)** Left side shows an overall view of the *Hodgkinia* LSU with the area where hL47 and hL44 are located highlighted. The middle and right shows zoomed-in views of h44/h47 in two representations: as molecular surfaces (center) and as cartoons with the secondary structure elements colored according to their genomic origin in exon 1, 2 (for hL44) or 2’ (for hL47; the genomic structure of this gene is detailed in **Figure S6**). (**B**) The interactions between exon 2’ of hL47 (shown as a cartoon) and hL44 (shown as surface potential or hydrophobicity surfaces). The h44/h47 interaction is driven by a cluster of hydrophobic phenylalanine residues present on exon 2’ of h47 that docks into a hydrophobic cavity on h44 (right), along with an ionic interaction between a lysine in hL47 and a negatively charged region of hL44 (middle).

### Three cicada proteins remodel the *Hodgkinia* central protuberance

The CP is an integral part of the LSU, directly interacting with the head of the small subunit and enabling the vectorial movement of tRNAs and mRNA to advance protein synthesis ^38^. The bacterial CP is composed of multiple bRPs and the 5S rRNA. The *Hodgkinia* CP is modified by the loss of bL31 and addition of hL38, hL46, and hL51, all of which interact with the 5S rRNA in distinct ways. While hL38 interacts with the 5S rRNA via stacking interactions (**Figure 3A**, 1-2), hL46 positions a short alpha-helix deep into the 5S rRNA, stabilizing its 5’-terminal end (**Figure 3A**, 4). It is worth noting that the N-terminal R26 residue is exactly the predicted sequence resulting from cleavage of the hL46 signal peptide (**Figure 3A**, 4; **Table S2**), confirming the computational signal sequence prediction of at least this protein. Finally, cationic resides of hL51 establish ionic interactions with the phosphate groups of the 5S rRNA backbone (**Figure 3A**, 3). Overall, the configuration of the *Hodgkinia* CP features a protein-rich environment compared to a normal bacterial CP, and results in a 5S rRNA almost entirely shielded from solvent (**Figure 3B**, left). While the addition of host proteins to mitoribosomes that still retain 5S rRNA has been reported, these proteins tend to bind to bRPs, not 5S rRNA ^8^. The burying of 5S rRNA in the *Hodgkinia* ribosome is perhaps most similar to the structural changes in the CP of eukaryotic cytosolic ribosomes (**Figure 3B**, right).

**Figure 3.**
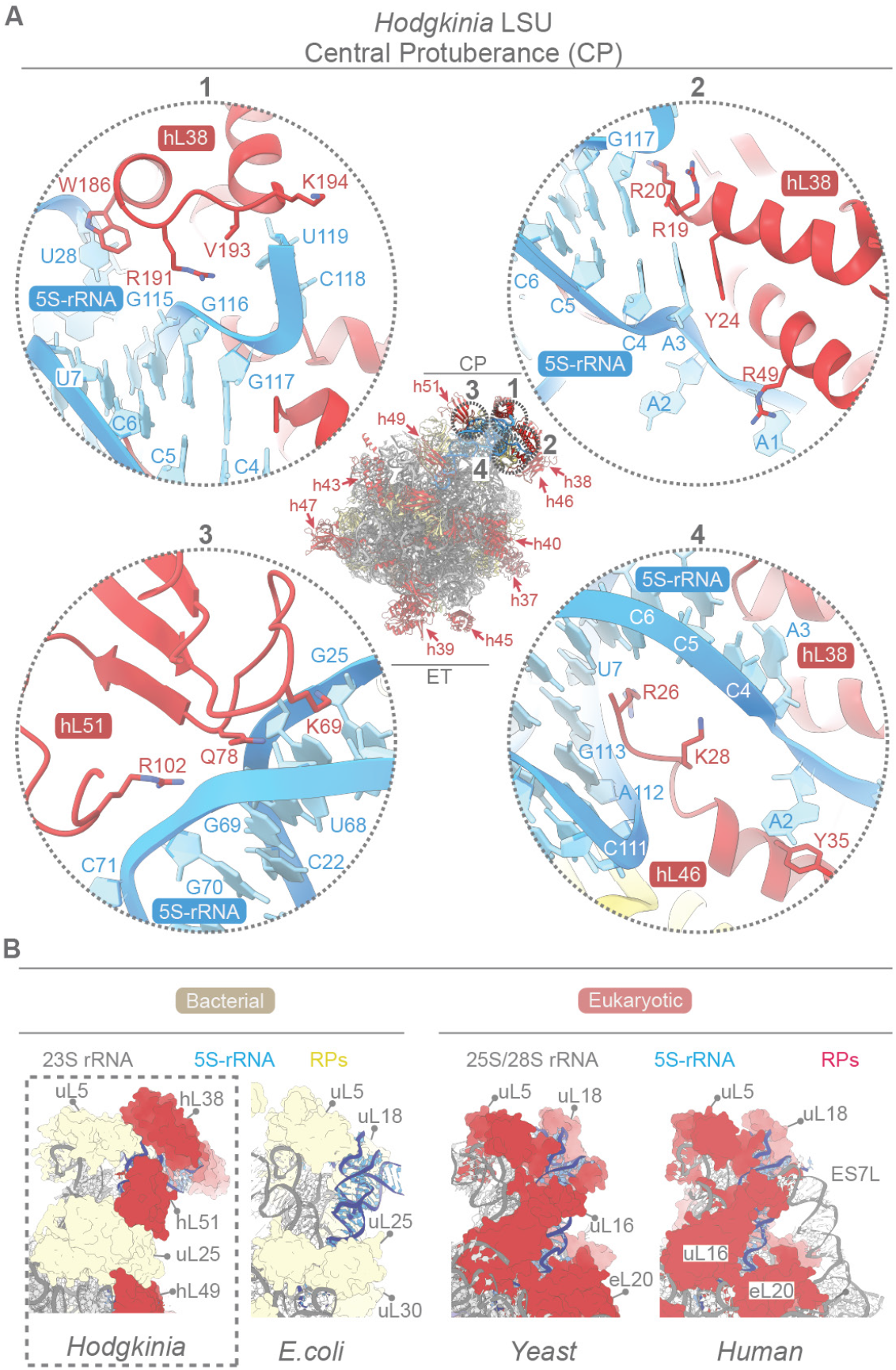
The central protuberance (CP) of the *Hodgkinia* LSU is highly modified. (**A**) Center, an overview of the *Hodgkinia* LSU with cicada proteins labeled in red. The CP and the exit tunnel (ET) areas are indicated. Regions 1, 2, 3, and 4 are zoomed-in views of the CP with cicada proteins colored red, *Hodgkinia* proteins of bacterial origin yellow, and the 5S rRNA blue. hL38 interacts with the 5S rRNA mainly by stacking interactions (1, 2) while hL51 (3) and hL46 (4) establish multiple ionic interactions with the 5S rRNA. (**B**) Comparative view of the CP from *Hodgkinia* (left, boxed), *E. coli*, and the *S. cerevisiae* and human cytosolic structures (right). The newly acquired cicada proteins almost fully shield the 5S rRNA from the solvent (left), in contrast to a canonical bacterial configuration, and more similar to the extensively covered 5S rRNA of eukaryotic cytosolic ribosomes (right).

### The addition of cicada proteins radically alters the outbound trajectory of the nascent peptide

The ET guides newly synthesized peptides towards the exterior of the LSU, and is thought to prevent premature collapse of the nascent peptide into non-native fold states ^39^. These functions are accomplished by architectural constraints imposed by bRPs and LSU rRNA residues upholstering the tunnel. The first half of the tunnel is straight and narrow, while towards the end it widens into a vestibule to allow the formation of secondary structure elements. Finally, the ET opens to a wide cavity where chaperones and other protein modifying enzymes can act on the nascent peptide. In most bacteria, the ET is comprised of LSU rRNA and six RPs: uL4, uL22, uL23, uL24, uL29, and bL32 ^39^. Of these six canonical ET-related ribosomal proteins, *Hodgkinia* retains only uL4 and uL22.

The *Hodgkinia* ET is most dramatically modified, however, by the addition of three cicada proteins (hL39, hL42 and hL50) at the end of the ET, which significantly alters the normal path of the nascent peptide towards the solvent (**Figure 4A-C**). Cicada protein hL39 directly blocks the trajectory of the exiting nascent chain (**Figure 4A**, right). Cicada proteins hL42 and hL50 are also located near the ET, but seem to play a peripheral role to orient and support hL39 (**Figure 4B**). We modeled the trajectory of the ribosome nascent chain (RNC) in our *Hodgkinia* map using a cryo-EM reconstruction from *E. coli* ribosomes bound to a P-site tRNA (PDB ID 7ZP8, **Figure 4D-E**). This analysis revealed that the position of cicada protein h39 imposes an almost 90° turn of the RNC trajectory compared to *E. coli* (**Figure 4D**).

**Figure 4.**
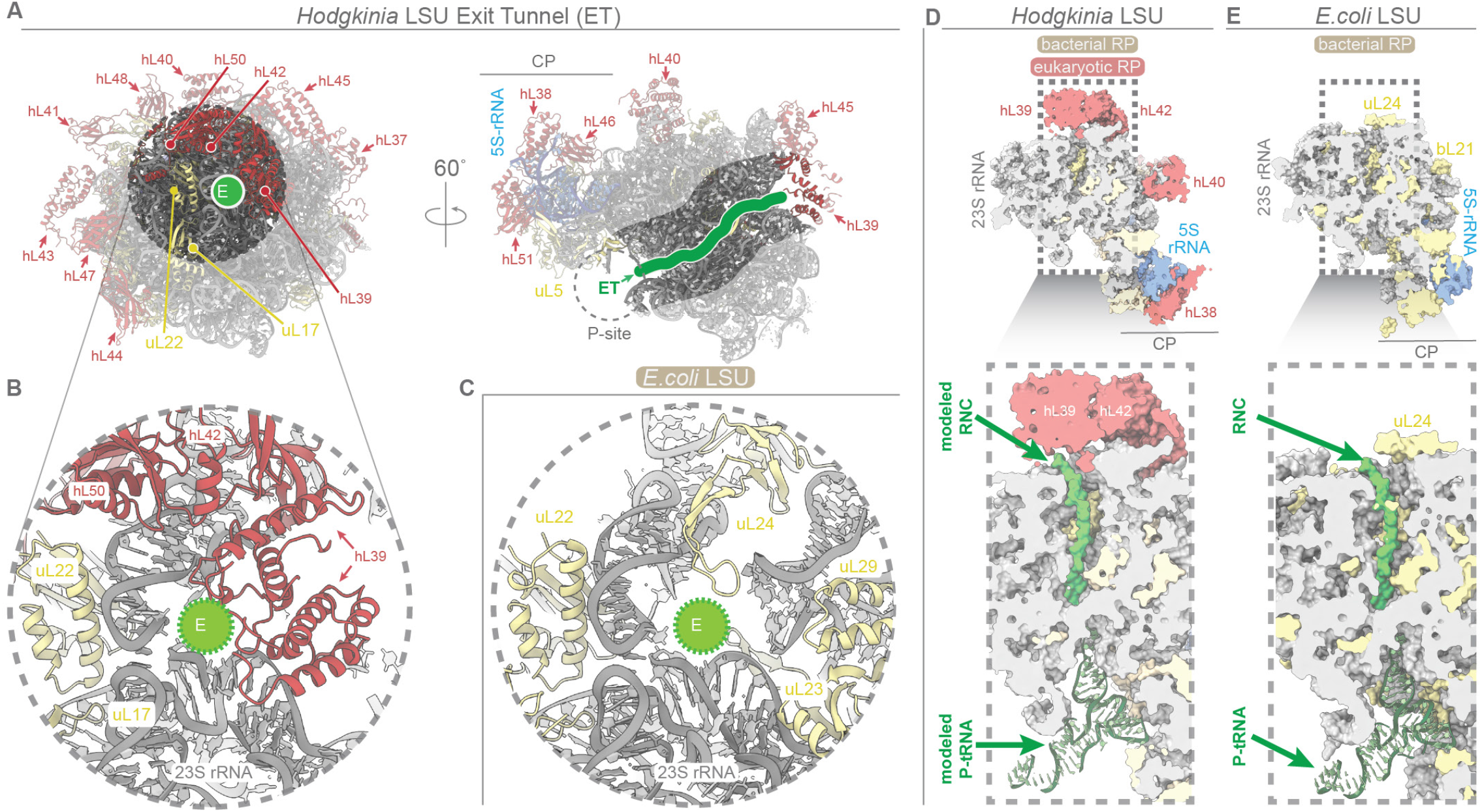
The exit tunnel (ET) of the *Hodgkinia* LSU is radically altered by a host cicada protein. (**A**) Two views of the *Hodgkinia* LSU, with a view looking straight into the ET (left) and one cut away for a side-on view (right). Ribosomal proteins from the cicada genome are labeled and colored red and ribosomal proteins of bacterial origin are colored yellow. The trajectory of the Ribosome Nascent Chain (RNC) is highlighted in green. (**B**) Zoomed-in view of the end of the ET, where cicada proteins hL39, hL42 and hL50 are located. (**C**) Equivalent view as in (B) from *E*.*coli* (PDB ID 7ZP8). Comparisons of the trajectory of the RNC in *Hodgkinia* (**D**) and *E. coli* (**E**), with 23S rRNA colored grey, 5S rRNA blue, ribosomal proteins of bacterial origin yellow, ribosomal proteins from the cicada red, and the peptidyl-tRNA in green. Cicada protein hL39 imposes a severe almost 90-degree turn of the RNC to access the solvent (D) compared to a typical bacterial ET (E).

## DISCUSSION

### The evolutionary context of the *Hodgkinia* ribosome

Comparative genomics suggests that the mitoribosome of the last eukaryotic common ancestor (LECA) retained 53 of the 54 standard alphaproteobacterial bRPs, having lost only bS20 ^1,3,4^. Genomic analyses also indicate that the LECA mitoribosome had acquired 24 mRPs (11 on the LSU, 13 on the SSU) before the divergence of the eukaryotic lineage ^1^. The *Hodgkinia* LSU has lost 9 bRPs and gained 15 new eukaryotic proteins, suggesting that it is in a more advanced state of protein loss but a similar level of protein gain compared to the inferred LECA LSU. The single structure we report here does not resolve whether *Hodgkinia* lost its bRPs before accumulating host proteins, or vice versa.

However, the large diversity of insect endosymbionts in various stages of genome reduction and host integration ^15^ might provide an avenue to resolve open questions related to when, how, and why new host proteins accumulate on endosymbiont (and organelle) ribosomes, and why some structural regions are more or less tolerant of change.

The relative order of mitoribosome rRNA reduction and expansion in relation to bRP loss and mRP accumulation has been debated ^8,9,40–43^, but it now seems clear that the accumulation of mRPs preceded most of the length variation observed in mitoribosome rRNA ^8,43,44^. Our results suggest that accumulation of host protein before alterations in rRNA length is a general feature of endosymbiotic bacteria and is not an evolutionary path exclusive to mitochondria. The length of *Hodgkinia*’s LSU rRNA is 2,767 nts, only 26 nts shorter and well within a standard deviation of the average length of a non-mitochondrial alphaproteobacterial LSU in the SILVA database (mean = 2,793 nts; stdev. = 161; n = 6,955) ^45^. While the length of LSU from *Hodgkinia*’s free-living ancestor is unknown, it seems unlikely that the accumulation of 15 nuclear-encoded hRPs was an adaptive response to a trivial change in rRNA length.

### Why do ribosomes from endosymbiotic bacteria lose bRPs and accumulate host proteins?

We have shown that the endosymbiont *Hodgkinia* has accumulated at least 15 cicada proteins that are now part of its ribosomal large subunit, and that these host proteins do not directly patch the holes made by the loss of bRPs (**Figures 1B** and **S3**). What role, then—if any—might these new host proteins play? It could be that some hRPs stabilize the ribosome at positions distant to the lost bRPs in an allosteric fashion by buffering the structural instability caused by bRP loss. It could be that the addition of hRPs at the *Hodgkinia* ET provides some sort of chaperone-like function for the wildly divergent (and likely unstable) *Hodgkinia* proteins as they exit the ribosome, similar to the role that trigger factor plays in the folding of nascent peptides leaving the ET ^46^. But it also could be that some of these host proteins accumulate for more subtle reasons, the origin of which might not initially be related to ribosome function at all. Several authors have suggested that constructive neutral evolution (CNE)-like mechanisms ^47^ might be responsible for at least some of the changes seen in mitoribosomes, and for the accumulation of complexity in the ribosome in general ^1,42–44,48–50^. Under CNE models, hRPs accumulate through transient interactions that serve no initial purpose, but become fixed into place by mutations in other parts of the ribosome that are only allowed in the presence of the new hRP. Because these ‘hidden’ mutations are deleterious in the absence of the nascent hRP ^51^, the interaction then becomes required for normal ribosome function ^49^.

The mitochondrion is the oldest endosymbiont known to science, and its origin was a key event in the evolution of eukaryotic life. But its antiquity, combined with a lack of comparative examples, has made understanding which features of mitochondria have arisen because of specific mitochondrial functions, and which have arisen as inevitable byproducts of the endosymbiotic process, difficult. Ribosomes from chloroplasts, which are the next oldest endosymbionts after mitochondria, also show alterations to the ET and accumulation of some eukaryotic proteins, but in general are much more bacteria-like in their structure ^52,53^. (It should be noted, however, that recent results suggest that this conservation of bacterial structure in chloroplast ribosomes might be due in part to a lack of sampling across archaeplastidal organisms ^54^.) Our results suggest that many of the striking features considered to be characteristic of mitoribosomes are likely to have arisen simply because the mitochondrion is a very old endosymbiont.

## ACKNOWLEDGEMENTS

We thank Bil Clemons and Jodi and Beta Puglisi for introducing I.S.F. to J.P.M.; Juan Alfonzo for introducing J.P.M. to Sergej Djuranovic; Sergej Djuranovic, Slavica Pavlovic-Djuranovic, and Jessey Erath for early access and assistance with the RAPPL protocol; and Jeremy Wideman, Bil Clemons, Ondřej Gahura, and Sergey Melnikov for comments on the manuscript. The cryo-EM data for this paper was collected on a ThermoFisher Titan Krios G2 cryo-transmission electron microscope funded by a National Science Foundation Major Research Instrumentation grant (NSF-MRI 1531991) and a Selectris X imaging filter and a Falcon 4i direct electron detector funded by a National Institutes of Health Shared Instrumentation Grant (SIG; 1S10OD036204-01). This work was supported by the Howard Hughes Medical Institute to J.P.M and J.M.M.; the National Aeronautics and Space Administration Interdisciplinary Consortia for Astrobiology Research (ICAR) grant number 80NSSC23K1357 to J.P.M.; and the Ministerio de Ciencia, Innovación y Universidades of Spain within the Plan Estatal de Investigación Científica, Técnica y de Innovación with grant number PID2023-147463NB-I00 to I.S.F.

## AUTHOR CONTRIBUTIONS

Conceptualization: J.P.M.; Methodology: I.S.F., J.P.M., J.M.W., J.N., N.S.; Investigation: all authors; Visualization: I.S.F., J.P.M., D.L.; Funding acquisition: J.P.M., I.S.F., J.M.W.; Project administration: J.P.M., I.S.F.; Supervision: J.P.M., I.S.F.; Writing – original draft: J.P.M., I.S.F.; Writing – review & editing: all authors.

## RESOURCE AVAILABILITY

The assembled *Diceroprocta semicincta* cicada genome has been deposited at DDBJ/ENA/GenBank under the accession JCCJVG000000000. The PacBio reads for the genome are available from the Sequence Read Archive under the accession SRR40353839, and the PacBio reads for the IsoSeq transcriptome analysis are available from the Sequence Read Archive under the accession SRR41210425. The genomic coordinates for the 15 hRPs have been deposited in GenBank with accession numbers QB309373-QB309387. Structural data has been deposited at the Protein Data Bank with ID 35CE.

## DECLARATION OF AI USE

During the preparation of this work, JPM used Claude Code to generate a custom Python script. This script was run with the following parameters (fasta_tax_ lengths.py SILVA_138.2_LSURef_tax_silva.fasta -t Alphaproteobacteria -x Mitochondria), and is available here: doi: 10.5281/zenodo.23423998. JPM also used Claude Science to organize the genome and transcriptome analyses for this work. JPM reviewed and edited the output as needed and takes full responsibility for the content of the published article.

## MATERIALS AND METHODS

### Ribosome purification

Two *Diceroprocta semicincta* cicadas were captured in Glendale, Arizona on 13 September 2025 and their abdomens were separated from their heads and thoraces. The heads and thoraces were placed in 70% ethanol and stored at −20°C for genome sequencing (see below). The bacteriomes were dissected from the abdomens in sterile phosphate-buffered saline, the buffer was removed, and the bacteriome tissue was flash frozen in liquid nitrogen and stored at −70°C. The ribosomes were purified using a slightly modified RAPPL protocol ^28^: frozen bacteriomes were thawed on ice for 2 min, resuspended in 500 µl of 4°C RAPPL lysis buffer, transferred to a 2 mL T-dounce and homogenized 15 times, and the mixture was then centrifuged at 20,000 X g for 20 min at 4°C. The remaining in-solution lysate was added to 30 µl RAPPL beads prewashed in RAPPL lysis buffer, mixed by gentle swirling, and incubated for 30 min at 4°C with gentle rocking. Beads were isolated in a magnetic stand and the solution was removed. The beads were washed 3X in 500 µl RAPPL wash buffer, and the ribosomes were released into 15 µl of RAPPL elution buffer for 15 mins at 4 °C with gentle rocking.

### Cryo-EM data acquisition

To visualize ribosomes via cryo-EM, 3 µL of the RAPPL-purified sample was applied to Quantifoil 2/1, 2 nm ultrathin carbon, 200-mesh gold TEM grids (Ted Pella, Inc., Redding, CA) after they were made hydrophilic by a 15 second glow discharge. Samples were allowed to adsorb on the grids for 2 minutes at 100% humidity before plunge freezing using a Vitrobot Mark IV and a liquid nitrogen cooled ethane slurry with 100% humidity at 16°C for 3 seconds blot time. Ribosome visualization was performed using a Krios G2 equipped with a Falcon4i/SelectrisX at 130,000 X nominal magnification, resulting in 27,282 EER movies with a 0.9308 Å/pixel size, an electron dose of 25 e^−^/Å^2^, and an energy filter slit width of 10 eV.

### Cryo-EM image processing

Data processing was entirely performed in Relion-5 ^55^ with CTF estimation in Ctffind4 ^56^. Eer files were motion corrected with the Relion implementation of the MotionCorr algorithm ^57^ using whole-frame alignment and a B-factor of 150 with sums of power spectra every 4e^−^/Å^2^. We used a FFT window of 1024 and the exhaustive search option turned on for ctf estimation in Ctffind4. We randomly selected a subset of 500 images for auto-picking with the Relion LoG picker using maximum diameter of 380Å and minimal of 120Å. We tried 3 different threshold parameters (0, 0.5 and 1) and manually checked the accuracy of the picking.

With a small set of particles originating from 500 images with the LoG picker, we performed 2 rounds of 2D classifications (T parameter 3 and 5) and manually selected those class averages that by shape and size looked like ribosomes. This curated small set of ~5,000 particles was used as a training set in Topaz ^58^. A first Topaz model was used to re-pick the set of initial 500 randomly selected images. After another round of two 2D classification jobs and manual selection of class averages reminiscent ribosomes, we trained a second Topaz model that significantly outperformed the first one in terms of number of particles and alternative views. We used the second Topaz model to pick the whole dataset, which generated an initial set of around 3 million particles. We then performed 5 iterative rounds of 2D classifications and manual selection of the 2D ribosome-looking class averages, subsequently increasing the T parameter at every 2D classification round. We used T=3 for the first round, T=4 for the second, T=5 for the third, T=6 for the fourth, and T=8 for the final fifth round. We did not attempt to separate ribosome populations at this stage, even though it was evident by the shape and details of the 2D class averages that there were multiple and very different ribosome populations in the sample. The final subset of particles selected after 2D classifications was in the range of 700,000. To separate the different ribosome populations we combined the Stochastic Gradient Descent (SGD) algorithm ^59^ and masked 3D classifications without alignment within Relion-5. Using the 700K dataset we launched an Initial Model job with 6 classes and T parameter of 2. The algorithm sorted ~70% of the particles to a class which we identified as the host insect cytoplasmic eukaryotic 80S ribosome. There was a second class of ~20% of the particles that resembled a prokaryotic large subunit and a small class that looked like a bacterial small subunit. We selected the particles of the class reminiscent of a bacterial large subunit and performed a 3D auto-refine job that generated a volume of good quality.

At this stage, we re-extracted the particles to a bigger box size (480 pixels) and performed another 3D refinement job that reached an overall resolution of 4 Å. We used Blush regularization for all 3D refinements ^60^. Next, after detailed analysis of the map, we performed a masked classification without alignment (4 classes, T=3). This classification job identified a homogenous subset of particles (53,191 particles) that refined to 3.5 Å resolution. A final iterative routine combining 3D auto-refinements with ctf-refinements ^61^ (defocus per-particle, astigmatism per micrograph) and Bayesian polishing improved the resolution to 3 Å ^62^. For model building we used Coot ^63^ with maps sharpened with Relion and phenix.autosharpen using the local sharpening option ^64^. The final model was stereochemically refined against the sharpened map in two steps: first, using phenix. real_space_refinement with secondary structure restraints ^65^ and after manually fixing multiple errors we performed a second refinement step in Fourier space using Refmac5 ^66^. The model statistics were analyzed with MolProbity ^67^.

### Protein identification in the cryo-EM map

After establishing that the map was from the *Hodgkinia* LSU, we identified the areas of unexplained density using ModelAngelo run with no input sequences (the build_no_ seq flag) followed by an HMM search (the hmm_search flag) against predicted cicada proteins. During most of the course of this work, we did not have access to the cicada genome, so we used a bacteriome-enriched transcriptome from the same cicada species (*Diceroprocta semicincta*) generated in 2010 for a different project ^26^. Putative proteins from these transcriptome data were generated with the LongOrfs package in TD2 ^68^ with the minimum protein length set to 40 and the requirements for canonical start and stop codons relaxed (the -m 40 --alt-start --all-stopless flags). While the ModelAngelo HMM search was initially performed against the 1.4 M proteins predicted from this method, and these data were used to identify the 15 cicada hRPs on the *Hodgkinia* LSU, we subsequently verified the presence and structure of these genes from a high-quality cicada genome and isoseq transcriptome generated in 2026, as described below.

Protein identifiers were extracted from the best_hits.csv file generated from the ModelAngelo HMM search against the TD2-predicted cicada proteome, and the protein sequences associated with these identifiers were used as input into ModelAngelo in an attempt to place these putative cicada proteins into the map. This process was iterative, as it was immediately clear that some of the cicada proteins were excellent fits in the map while others were less clear, especially in regions of the map that were of lower quality. AlphaFold 3 was used to predict the structure of the identified proteins and, in general, the predictions featured a central core of secondary structure-rich elements along with extended N- and C-termini with unstructured tails. We roughly placed well-predicted central core regions of the

AlphaFold 3 models in the map, using the short but well-defined stretches built by ModelAngelo as a guide. Many of the cicada proteins showed well defined N- and C-termini in the map, usually stabilized by molecular contacts with adjacent ribosomal proteins. These fragments were built manually.

### Cicada genome analysis

DNA for genome sequencing was isolated from small amounts of cicada tissue from the samples stored in ethanol at −20°C on 13 September 2025 (the same insects used to solve the *Hodgkinia* LSU structure, as described above). DNA was extracted using the Nanobind PanDNA (PacBio) high molecular weight DNA extraction kit, following the additional “Ethanol-preserved tissues” pretreatment outlined in the Nanobind PanDNA kit instructions. Extracted DNA was sent to SeqCenter (Pittsburgh, PA) for sequencing. Genomic DNA was fragmented to 40-50 kb using the Megaruptor 3 instrument from Diagenode, and sequenced on a PacBio Revio. PacBio sequencing produced 5.6 M reads with an average length of 16 kb for a total output of 88.5 Gb. These reads were assembled using hifiasm ^69^ using the default purge level (-l 2), which produced an assembly of 4.9 Gb in 1,396 contigs with an N50 of 13.2 Mb and an average coverage of ~18X. This 4.9 Gb genome size is in line with other cicada genomes on GenBank (4.45–6.52 Gb; GCA_979322825.1, GCA_980043705.1, GCA_034769895.1), and the BUSCO v.6.0.0 completeness was 98.6% when using the hemiptera_odb10 lineage dataset of 2,510 BUSCOs ^70^, suggesting that the genome assembly was of sufficient quality for gene finding and the verification of hL37-hL51 on the cicada genome.

### Transcriptome sequencing and genome annotation

The bacteriomes from two cicadas caught in July 2026 were flash frozen in liquid nitrogen. Samples were thawed on ice and total RNA was extracted with TRIzol using manufacturer’s protocol. Total RNA was frozen at −70° C and shipped shipped to SeqCenter (Pittsburgh, PA) for IsoSeq v2 sequencing on a PacBio Revio instrument. 22,768,079 reads were generated from two bacteriome libraries. To find reads encoding the 15 cicada proteins identified from the cryo-EM map, the IsoSeq reads were translated using TD2.LongOrfs (minimum length set to 50aa), which yielded 78,852,357 candidate peptides. The 15 proteins were searched against these predicted peptides using BLAST+ v2.12.0: blastp -evalue 1e-10 -max_target_ seqs 1000000, which identified 7,691 reads as containing the 15 proteins with ≥ 90% of that protein’s length with 5 or fewer amino acid substitutions. For each protein, a consensus sequence was built from the matching reads by taking the majority residue at each position across all read-containing ORFs, aligned in protein coordinate space. These 15 read-derived consensus sequences were aligned to the *D. semicincta* genome using miniprot v0.18: miniprot --gff -G 200k, which defined the CDS coordinates for each locus. Separately, all 22,768,079 IsoSeq reads were mapped to the cicada genome using minimap2 v2.28: minimap2 -ax splice:hq -uf -I 8G -G 500k. For each of the 15 loci, reads were counted if their alignment spanned the full CDS with no splice junctions outside the CDS model. This structural criterion, applied without reference to the protein sequence, identified 8,004 supporting reads across the 15 loci. Some additional reads were recruited in this step due to some protein predictions falling outside the 90% length cutoff described in the previous paragraph.

### *Hodgkinia* LSU rRNA length analysis

LSU rRNA sequence lengths were analyzed using the SILVA_138.2_LSURef_tax_silva.fasta file downloaded from the SILVA database ^45^ on 2 June 2026. Mean, median, and standard deviation of 6,955 alphaproteobacterial LSU sequences were computed after excluding mitochondrial sequences as well as those derived from metagenomic sampling.

---

**Figure S1.**
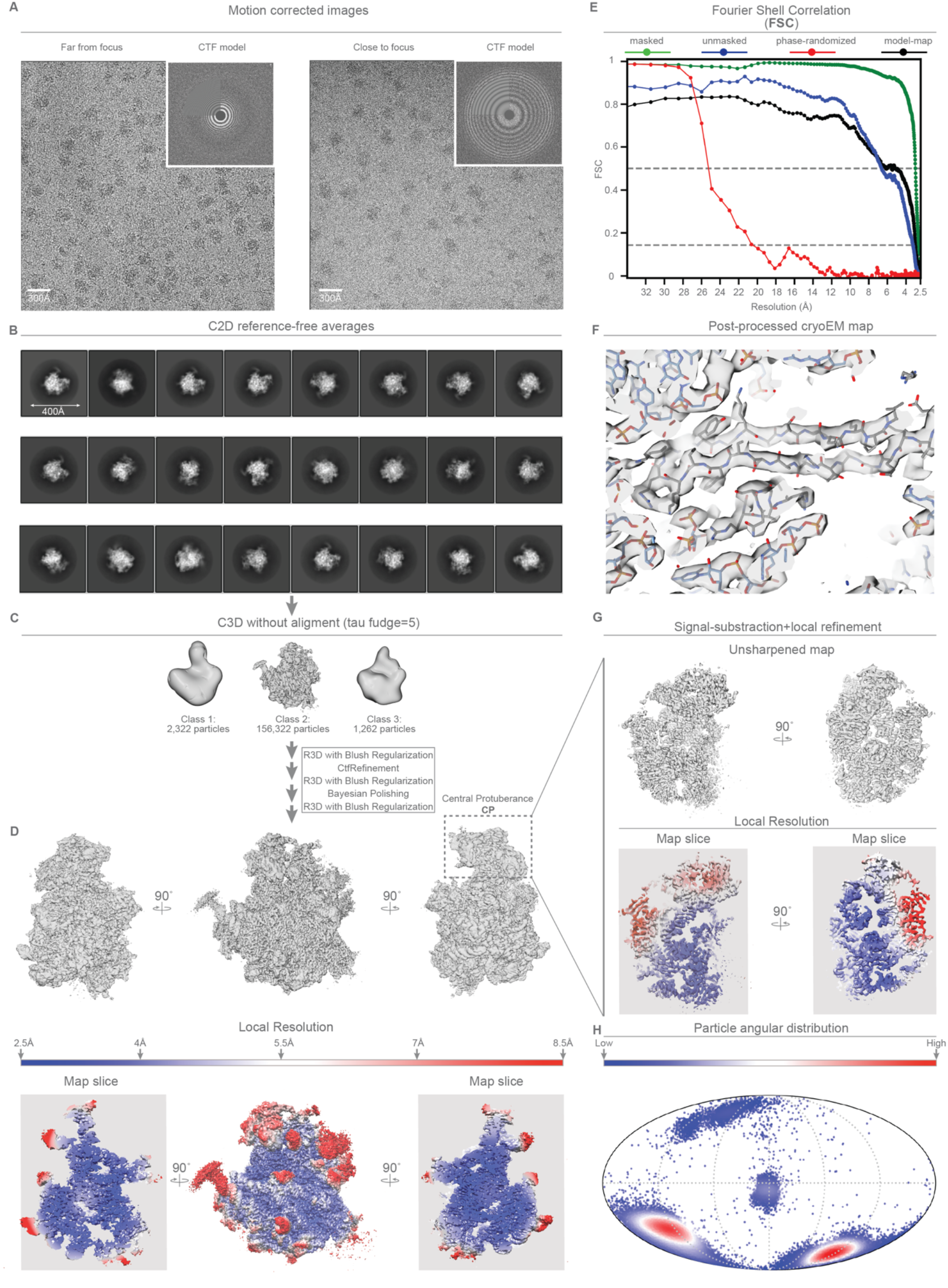
(**A**) Representative motion corrected images of the dataset. (**B**) 2D class averages selected for 3D refinements and classifications. (**C**) Simplified schematics of the data processing workflow followed to reach the final map. (**D**) Top, orthogonal views of the final unsharpened maps of the *Hodgkinia* LSU. The bottom shows the same map colored according to local resolution estimations. (**E**) The overall resolution was estimated to be 3Å. Regions at the core of the particle like the peptidyl transferase center (PTC) exhibited map features compatible with resolutions better than 3 Å, however the quality of the map degraded towards the periphery with local resolution estimation in the 4-5 Å range. (**F**) Post-processing using a global B-factor of −20 Å^2^ produced a well-defined map. (**G**) The central protuberance (CP) region was independently refined using local signal subtraction and refinement, producing and enhanced map that allowed better modeling. (**H**) The angular particle distribution for the final refinement job using a Mollweide projection of the spherical values.

**Figure S2.**
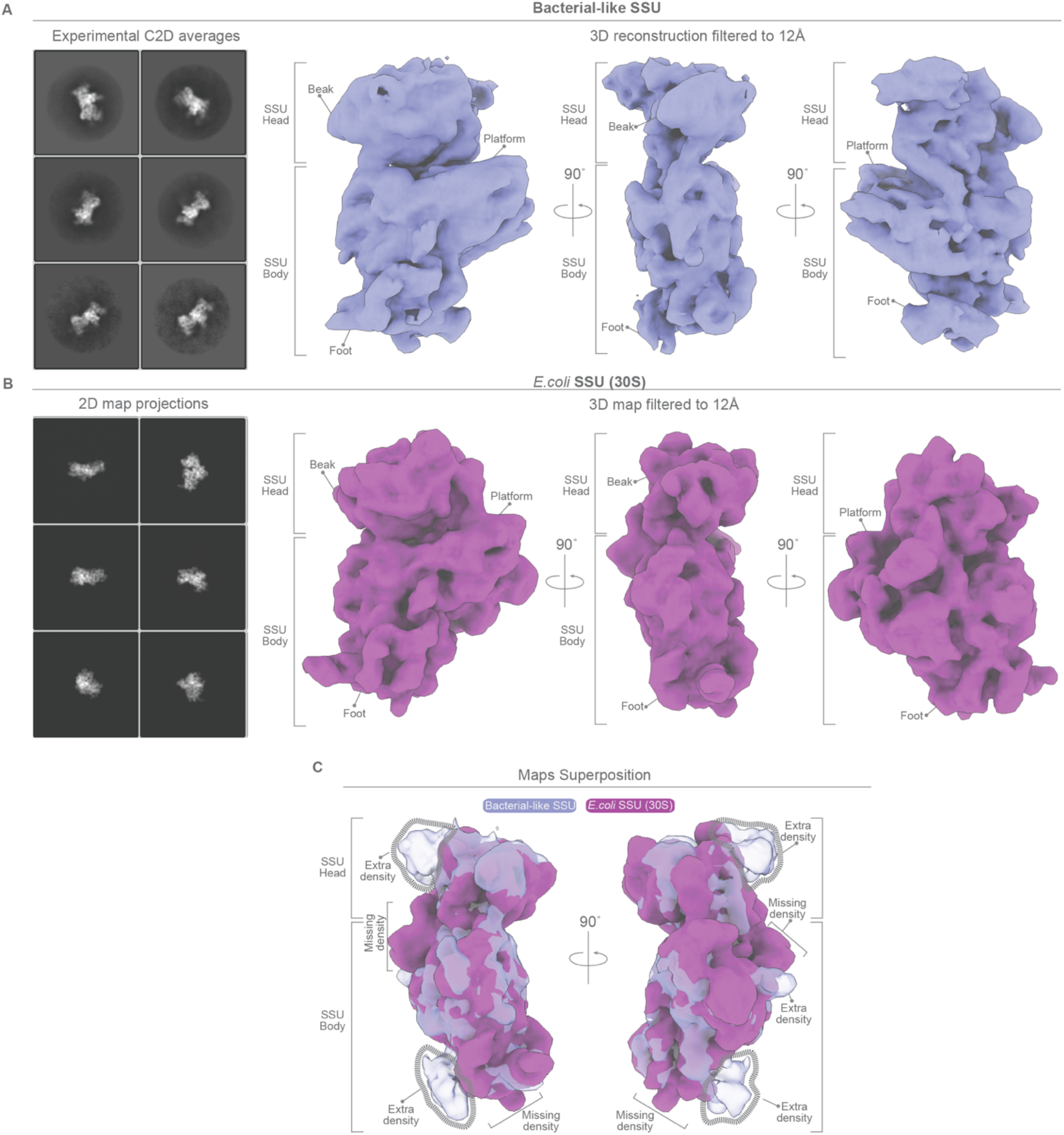
(**A**) 2D class averages from the dataset that resemble a prokaryotic small subunit (SSU). This set of particles was small (~2,500), however it was enough to produce a 3D map with a resolution of around 12Å. This map is shown in 3 different orthogonal views with the characteristic regions of the SSU indicated. (**B**) For comparative purposes, we filtered a SSU map from *E. coli* to 12Å and projected that map in 2D to generate a set of views in similar orientations as our experimentally identified 2D class averages. On the right, we show the *E*.*coli* map filtered to 12Å in similar orientations as in (A). (**C**) When we superpose these two maps, two large regions of extra density are evident at the top of the head and at the foot of the SSU. At this resolution we are unable to identify this SSU as from *Hodgkinia*, or identify proteins that make up this missing density.

**Figure S3.**
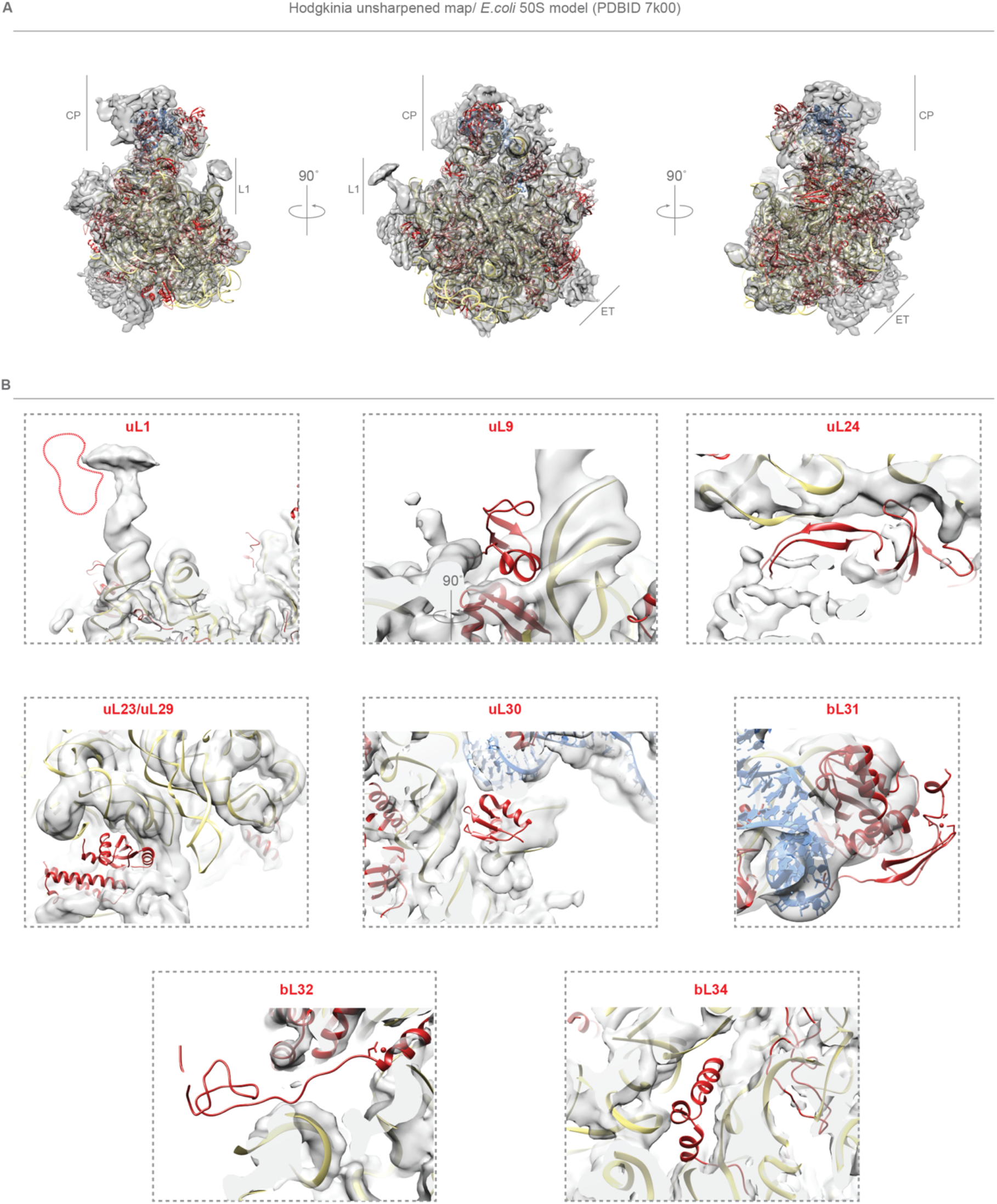
(A) *Hodgkinia’s* genome suggested that several ribosomal proteins (RPs) might be missing, as well as extra unexplained density compared to *E. coli* (gray, unfilled areas of the map). We fitted a whole *E*.*coli* model (PDBID 7k00) into the unsharpened map, using the highly conserve peptidyl transferase center (PTC) region of the ribosomal 23S rRNA as an initial anchor and superposed the rest of the *E*.*coli* LSU afterwards. (B) We then looked for the positions on the *E*.*coli* LSU of the proteins predicted to be missing in *Hodgkinia* (uL1, bL9, bL21, uL23, uL24, uL29, uL30, bL31, bL32 and bL34) and found clear evidence of missing density for all but bL21. Based on the solid cryo-EM density for bL21 we went back to the genome and found an annotation error, which we confirmed and fixed.

**Figure S4.**
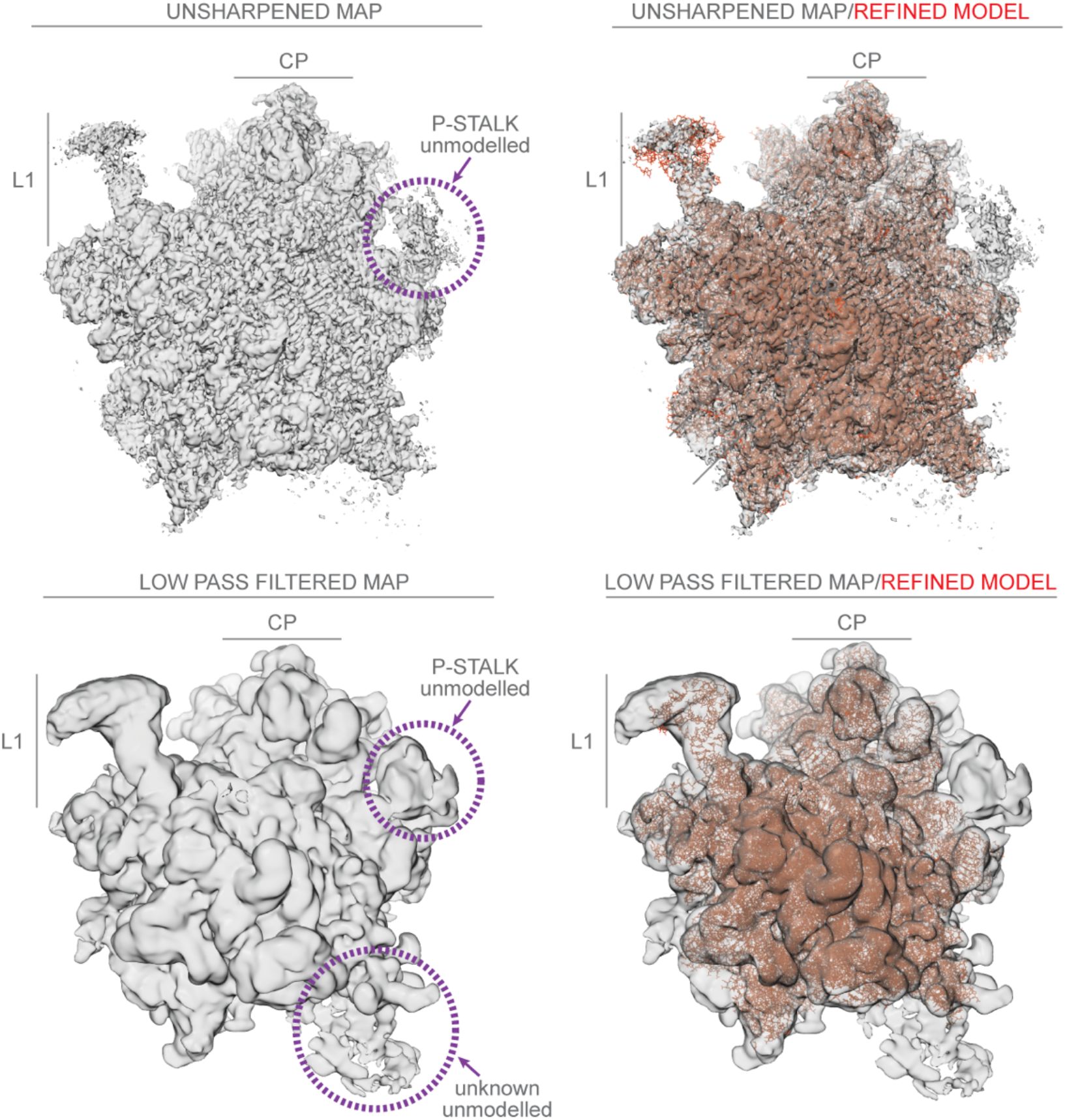
Top left, crown view of the final unsharpened map of the *Hodgkinia* LSU with the L1 stalk, the central protuberance (CP) and the acidic (P) stalk indicated. Due to the mobility of the P-stalk we were unable to build the corresponding atomic model. On the top right we show the unsharpened final map of the *Hodgkinia* LSU as semi-transparent cryo-EM density with the final stereochemically refined model inserted with red sticks representation. The majority of the map was interpreted in atomic terms, however, low-pass filtering of this map (bottom row) revealed additional peripheral densities which we were unable to identify and thus were left unmodeled.

**Figure S5.**
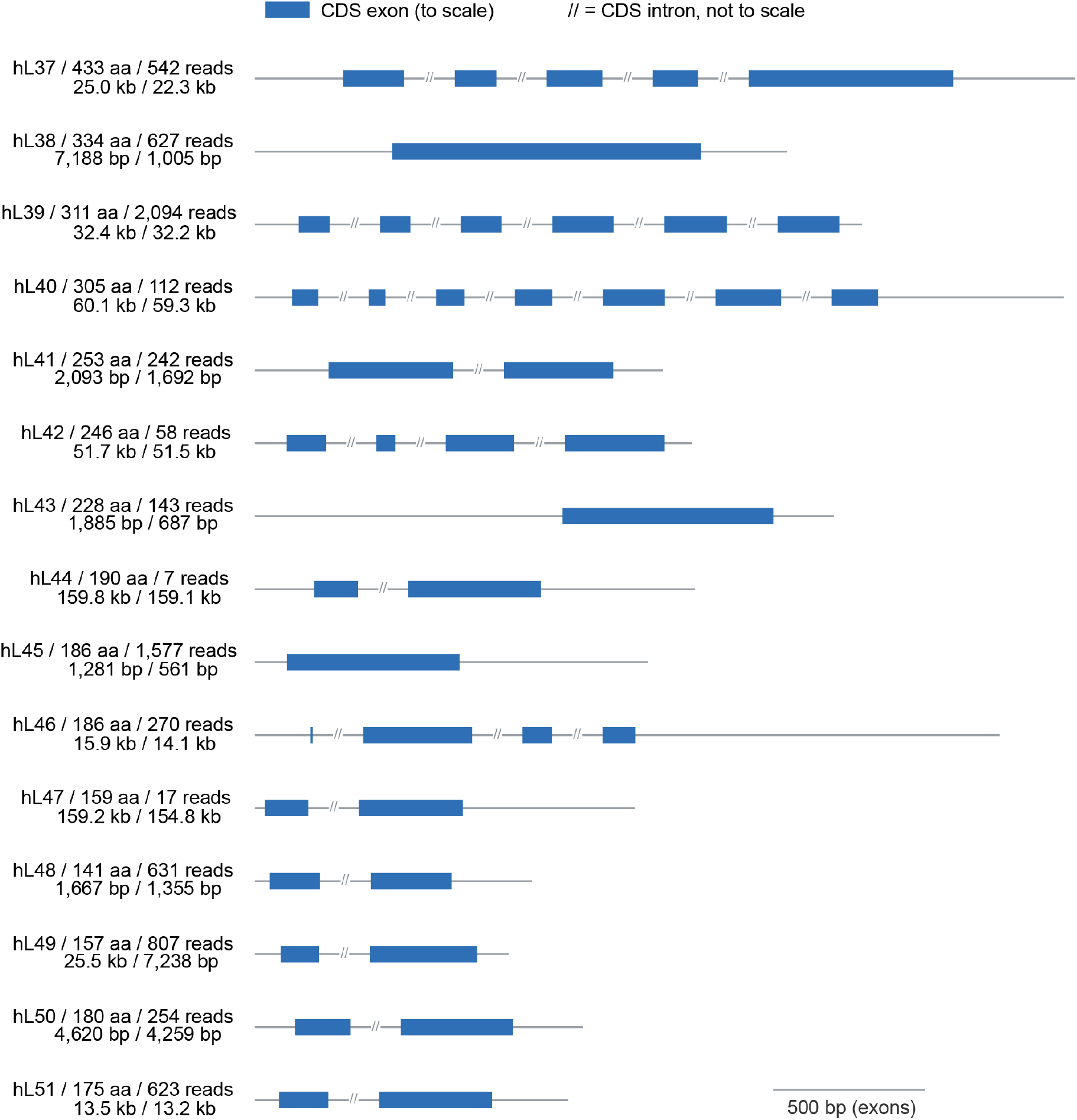
Genomic structure of the genes for the 15 host proteins on the *Hodgkinia* ribosome. On the left, each gene is labeled with the protein name / length in amino acids / number of IsoSeq reads covering this exact splice form on the top line, and the length of the complete gene including the predicted UTRs / length of the coding sequence (CDS) on the bottom line. Exons are drawn to scale, while the introns are reduced in size by hash marks (//) and are not drawn to scale.

**Figure S6.**
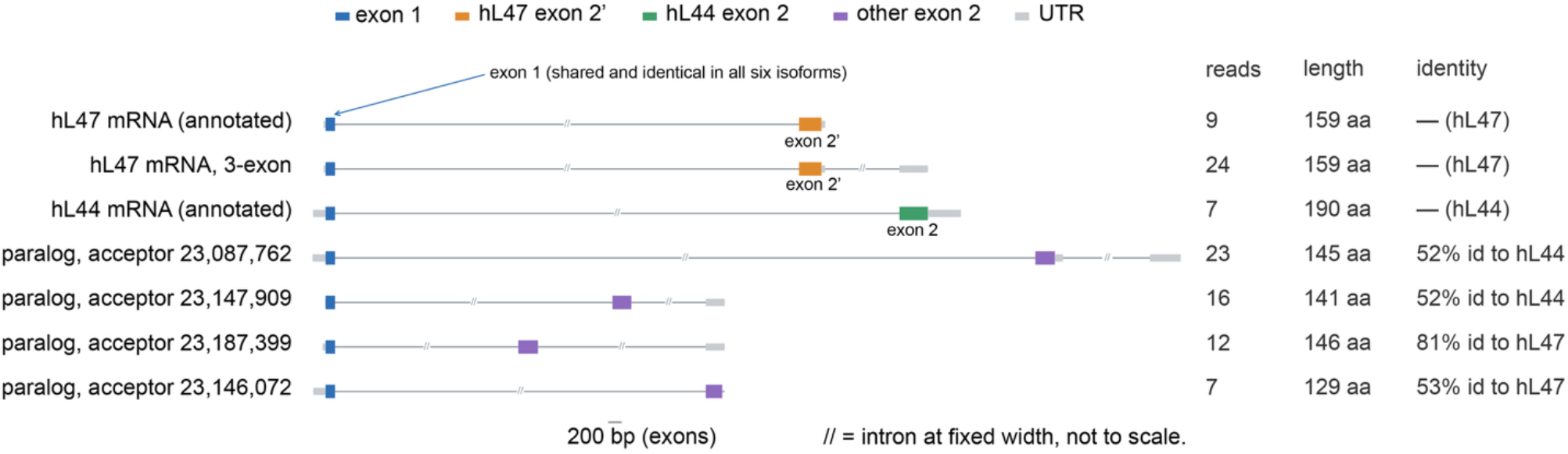
Splice isoforms for the cicada gene that produces hL44 and hL47. All six predicted isoforms share the same exon 1 (blue) on contig ptg000184l (coordinates for hL44 and hL47 are in Table S2). The cicada protein hL44 is produced by a unique exon 2 (green), while hL47 is produced by exon 2’ (orange). Two different versions of the hL47 transcript exist (top two rows) but they produce exactly the same protein product so it is unclear which transcript produces the hL47 protein, or if both do. Exon lengths are to scale but intron lengths are not, although the relative sizes of the introns are representative. The introns for this gene are large: the distance between exon 1 and exon 2’ in hL47 is 154 kb, and the distance between exon 1 and exon 2 in hL44 is 158 kb.

**Table S1.** Cryo-EM data collection, refinement, and validation statistics.

|  | *Hodgkinia* LSU<br>(EMD-100351) |
|---|---|
| **Data collection and processing** |  |
| Magnification | 130,000x |
| Voltage (kV) | 300 |
| Electron exposure (e–/Å<sup>2</sup>) | 25.93 |
| Defocus range (μm) | -0.8 to -2.5 |
| Pixel size (Å) | 0.93 |
| Symmetry imposed | C1 |
| Initial particle images (no.) | 3224334 |
| Final particle images (no.) | 156322 |
| Map resolution (Å) | 3 |
| FSC threshold | 0.143 |
| **Refinement** |  |
| Initial model used (PDB code) | ModelAngelo/7k00<br>Manual build |
| Model resolution (Å) | 3.5 |
| FSC threshold | 0.5 |
| Model resolution range (Å) | 3 |
| Map sharpening *B* factor (Å<sup>2</sup>) | -20 |
| Model composition |  |
| Non-hydrogen atoms | 95,574 |
| Protein residues | 4,789 |
| Nucleotide residues | 2,652 |
| *B* factors (Å<sup>2</sup>) |  |
| Protein | 94.35 |
| Nucleotide | 88.56 |
| R.m.s. deviations |  |
| Bond lengths (Å) | 0.0091 |
| Bond angles (°) | 1.214 |
| Validation |  |
| MolProbity score | 2.06 |
| Clashscore | 4.75 |
| Poor rotamers (%) | 1.67 |
| Ramachandran plot |  |
| Favored (%) | 85.9 |
| Allowed (%) | 12.51 |
| Disallowed (%) | 1.75 |

**Table S2.** Predicted characteristics, annotation, and genome coordinates for the fifteen cicada proteins on the *Hodgkinia* LSU. The signal peptide cleavage sites were predicted using SignalP 6.0 ^S1^. The consensus annotation was made by combining the InterProScan ^S2^ results with BLASTP ^S3^ searches against the nr database, along with Foldseek ^S4^ searches using AlphaFold 3 ^35^ structure predictions as search seeds.

| Name | Contig | Strand | CDS start | CDS stop | Len. (aa) | Signal peptide | Consensus annotation | InterPro evidence | BLASTP evidence | Foldseek evidence |
|---|---|---|---|---|---|---|---|---|---|---|
| hL37 | ptg000103l | + | 2601317 | 2623596 | 433 | Y, 17 and 18 | Gamma-interferon-inducible lysosomal thiol reductase (GILT) | PANTHER PTHR13234 (aa 56-263, e=5.1e-29) | GILT-like protein 1 isoform X2 (acc XP_039286794.1, e=6.73e-26) | Gamma-interferon-inducible lysosomal thiol reductase-like protein (AFDB Q7PQD1, prob=1.00, e=5.00e-17) |
| hL38 | ptg000311l | - | 35669526 | 35668522 | 334 | Y, 17 and 18 | Uncharacterized | No hit | hypothetical protein (KAG8297639.1, e=3.05e-04, 22%) | No hit |
| hL39 | ptg000062l | - | 3944712 | 3912484 | 311 | Y, 29 and 30 | UMP-CMP kinase 2, mitochondrial-like | No hit | UMP-CMP kinase 2 (KAE8746981.1, e=1.77e-10) | UMP-CMP kinase 2, mitochondrial (AFDB Q5EBM0, prob=1.00, e=1.24e-9) |
| hL40 | ptg000062l | + | 3677654 | 3736996 | 305 | N | UMP-CMP kinase 2, mitochondrial-like | GENE3D G3DSA:3.40.50.300 (aa 104-294, e=6.1e-08) | UMP-CMP kinase 2 (XP_021920734.1, e=3.99e-31) | UMP-CMP kinase 2, mitochondrial (AFDB Q5EBM0, prob=1.00, e=5.48e-13) |
| hL41 | ptg000184l | + | 50547705 | 50549396 | 253 | Y, 27 and 28 | Beta-1,3-glucan-binding protein (BGBP) | PFAM PF15886 (aa 121-231, e=1.1e-09) | hypothetical protein (acc KAJ0178108.1, e=4.37e-07) | Beta-1,3-glucan-binding protein (Fragment) (AFDB O96363, prob=1.00, e=3.56e-7) |
| hL42 | ptg000131l | - | 36660454 | 36608972 | 246 | Y, 20 and 21 | Magnesium-dependent phosphatase 1 (MDP-1) | PFAM PF12689 (aa 34-191, e=4.6e-31) | magnesium-dependent phosphatase 1-like (XP_049784562.1, e=8.03e-29) | Magnesium-dependent phosphatase 1 (AFDB Q86V88, prob=1.00, e=1.30e-17) |
| hL43 | ptg000036l | + | 9899134 | 9899820 | 228 | Y, 42 and 43 | Uncharacterized | No hit | No hits | No hit |
| hL44 | ptg000184l | - | 23277937 | 23118821 | 190 | Y, 17 and 18 | Beta-1,3-glucan-binding protein (BGBP) | PFAM PF15886 (aa 31-130, e=1.0e-02) | No hits | Beta-1,3-glucan-binding protein (Fragment) (AFDB O96363, prob=1.00, e=4.13e-6) |
| hL45 | ptg000311l | + | 35435798 | 35436358 | 186 | Y, 24 and 25 | Uncharacterized | No hit | No hits | No hit |
| hL46 | ptg000219l | - | 11203321 | 11189178 | 186 | Y, 25 and 26 | Reeler/ferric-chelate-reductase-like fold | No hit | No hits | Putative defense protein 3 (AFDB Q0Q028, prob=1.00, e=6.73e-7) |
| hL47 | ptg000184l | - | 23277937 | 23123102 | 159 | Y, 17 and 18 | Beta-1,3-glucan-binding protein (BGBP) | PFAM PF15886 (aa 31-117, e=7.2e-03) | unnamed protein product (acc CAG9861070.1, e=1.90e-02) | Beta-1,3-glucan-binding protein (Fragment) (AFDB O96363, prob=1.00, e=1.10e-5) |
| hL48 | ptg000184l | - | 28871256 | 28869902 | 141 | Y, 18 and 19 | Uncharacterized | No hit | No hits | No hit |
| hL49 | ptg000311l | + | 588695 | 595932 | 157 | Y, 26 and 27 | MBF2 / probable salivary-secreted-peptide-like | PFAM PF15868 (aa 34-125, e=2.3e-12) | probable salivary secreted peptide (XP_069694665.1, e=1.34e-04) | Probable salivary secreted peptide (AFDB D8KY57, prob=1.00, e=2.29e-9) |
| hL50 | ptg000184l | - | 30712044 | 30707786 | 180 | Y, 23 and 24 | Uncharacterized | No hit | No hits | No hit |
| hL51 | ptg000184l | + | 29426953 | 29440152 | 175 | Y, 16 and 17 | Beta-1,3-glucan-binding protein / GNBP family | PFAM PF15886 (aa 36-120, e=1.7e-03) | No hits | Gram-negative bacteria-binding protein 3 (AFDB Q9NHA8, prob=1.00, e=4.2e-06) |

